# Fatty acid profiles in adipose tissues and liver differ between horses and ponies

**DOI:** 10.1101/463018

**Authors:** Stephanie Adolph, Carola Schedlbauer, Dominique Blaue, Axel Schöniger, Claudia Gittel, Walter Brehm, Herbert Fuhrmann, Ingrid Vervuert

## Abstract

Fatty acids, as key components of cellular membranes and complex lipids, may play a central role in endocrine signalling and the function of adipose tissue and liver. Thus, the lipid fatty acid composition may play a role in health and disease status in the equine. This study aimed to investigate the fatty acid composition of different tissues and liver lipid classes by comparing Warmblood horses and Shetland ponies under defined conditions. We hypothesized that ponies show different lipid patterns than horses in adipose tissue, liver and plasma. Six Warmblood horses and six Shetland ponies were housed and fed under identical conditions. Tissue and blood sampling were performed following a standardized protocol. A one-step lipid extraction, methylation and trans-esterification method with subsequent gas chromatography was used to analyse the total lipid content and fatty acid profile of retroperitoneal, mesocolon and subcutaneous adipose tissue, liver and plasma. In the adipose tissues, saturated fatty acids (SFAs) and n-9 monounsaturated fatty acids (n-9 MUFAs) were most present in ponies and horses. N-6 polyunsaturated fatty acids (n-6 PUFAs), followed by SFAs, were most frequently found in liver tissue and plasma in all animals. Horses, in comparison to ponies, had significantly higher n-6 PUFA levels in all tissues and plasma. In liver tissue, horses had significantly lower hepatic iso-branched-chain fatty acids (iso-BCFAs) than ponies. The hepatic fatty acid composition of selected lipid classes was different between horses and ponies. In the polar PL fraction, horses had low n-9 MUFA and n-3 PUFA contents but higher n-6 PUFA contents than ponies. Furthermore, iso-BCFAs are absent in several hepatic lipid fractions of horses but not ponies. The differences in fatty acid lipid classes between horses and ponies provide key information on the species- and location-specific regulation of FA metabolism, thus affecting health and disease risk.

## Introduction

The physiological fundamentals of the lipid metabolism of equids are poorly understood. Several studies have shown that lipid and lipoprotein statuses differ among horse breeds [1-5]. Ponies have higher plasma lipoprotein contents than horses and seem to be more susceptible to developing hyperlipidaemia under a negative energy balance [2,5,6]. However, little is known about the impact of fatty acid (FA) profiles on lipid metabolism and homeostasis in different horse breeds. FAs are integral parts of cellular membranes and complex lipids such as triacylglycerides (TAGs) and phospholipids (PLs). They are involved in various general and specific biological processes that act to regulate cell and tissue metabolism, function and cellular signalling, thus affecting health, welfare and disease risk [7,8]. Specifically, polyunsaturated FAs (PUFAs) of the n-3 and n-6 FA family seem to be metabolically related to health conditions and inflammation. PUFAs of the n-3 series rather than n-6 PUFAs have commonly been shown to exert molecular actions that result in an improved risk factor profile in relation to metabolic and inflammatory dysregulations [9-11]. It is further speculated that the health impact of n-3 PUFAs on whole body homeostasis is mediated by resetting the adipose tissue (AT) function [12]. AT is no longer considered a simple fat storing tissue but rather contributes as an integrative key regulator in energy homeostasis and systemic metabolism [12,13]. AT can influence and communicate with many other tissues, including the brain, heart, vasculature, muscle and liver, on different molecular levels by releasing pro- and anti-inflammatory mediators such as interleukin 1 beta (IL-1β), interleukin 6 (IL-6), tumour necrosis factor alpha (TNF-α) and other adipokines [13,14]. In equids, a close link between AT function and health conditions is postulated [15]. Controversial studies about whether visceral fat or subcutaneous fat depots have a greater modulating impact on inflammation exist [16,17]. Thus, it is important to address how tissues vary with respect to FA composition in horses and ponies.

The aim of the current study was to compare the FA contents and profiles of different ATs, liver, plasma, and hepatic lipid classes between Shetland ponies and Warmblood horses. Considering the differences in the lipoprotein metabolism of horses and ponies, it was hypothesized that the FA profiles, with a special focus on the n-6 and n-3 PUFA dynamics, were different between equine breeds.

## Materials and methods

### Animals and preselection criteria

Six Shetland ponies with a mean age (± SD) of 6 ± 3 years and six Warmblood horses with a mean age (± SD) of 10 ± 3 years, all geldings with a median body condition score of (25^th^ /75^th^ percentile) 3.7 (2.2/4.4) for ponies and 3.6 (3.1/4.2) for horses on a scale of 1 to 6 [18] and a mean body weight (BW) (± SD) of 118 ± 29 kg (ponies) and 589 ± 58 kg (horses), were included in the study. Animals were individually housed in box stalls and bedded on straw. All animals had turnout onto a sand paddock for at least 5 h a day. During a two-week acclimatization period to the experimental procedure, animals were fed daily 2 kg meadow hay/100 kg BW, which was divided into two equal portions, one offered in the morning and one offered in the evening.

The animals had free access to water at all times. All animals were assessed for plasma adrenocorticotropic hormone (ACTH) to rule out pituitary pars intermedia dysfunction (PPID). Insulin dysregulation was excluded for all animals, as fasting serum insulin values were under the threshold of < 20 μU/ml after the combined glucose-insulin test (CGIT) performed according to Eiler et al. [19]. The project was approved by the ethics committee for animal rights protection of the Leipzig district government, in accordance with German legislation for animal rights and welfare (No. TVV 32/15). The study was part of a larger project on the consequences of increasing BW gain in horses and ponies.

### Blood collection

Aliquots of blood samples were collected by jugular vein puncture into tubes coated with sodium fluoride or a coagulation activator after an 8 h overnight fast to determine plasma glucose, serum insulin and plasma lipid FA composition. Samples were allowed to clot for 30-60 min before centrifugation. Plasma and serum were removed and stored at −80 °C until analysis.

### Adipose and liver tissue collection

For tissue collection, animals were sedated with romifidine (0.04 mg/kg BW, Sedivet®) and butorphanol (0.03 mg/kg BW, Alvegesic®). General anaesthesia was induced with 0.08 mg/kg BW diazepam (Diazepam-Lipuro®) and ketamine (3 mg/kg BW, Ursotamin®). Animals were orotracheally intubated, and anaesthesia was maintained with isoflurane (Isofluran® CP). Animals were placed in dorsal recumbency on a padded surgical table. After aseptic preparation, a 20 cm ventral midline incision was performed from cranial to the umbilicus. AT (~5 g at each location) was collected from the margins of incision (retroperitoneal (RPN)) and the mesocolon (MSC) of the descending colon (= visceral fat). Liver tissue (~2 g) was collected by biopsy forceps. After the abdomen was closed, the animals were repositioned in lateral recumbency. After aseptic skin preparation, incisions (~4 cm) were performed lateral to the tail head and in the middle of the neck at the nuchal crest. Approximately 5 g AT (subcutaneous (SC)) was collected from each location. A portion of each tissue biopsy specimen was stored in formalin, and the remainder was immediately flash frozen and stored in liquid nitrogen until analysis. After the procedure, horses and ponies were placed in a well-padded box to recover from general anaesthesia. All animals were treated with 1.1 mg/kg BW flunixin-meglumine (Flunidol® RPS 50 mg/ml) for three days.

### Plasma glucose and serum insulin

Plasma glucose concentrations were determined using the glucose oxidase (GOD)/peroxidase (POD) method [20]. Serum insulin was analysed by an immunoradiometric assay (IRMA)(^125^I) kit for human insulin (0-500 μIU/ml (0-17.5 ng/ml) (Demeditec Diagnostics GmbH, Kiel, Germany).

### Total FA profile and lipid class FA composition

#### Thin-layer chromatography

The liver (0.1 g) and fat samples (0.5 g) were cut into small pieces and put into 10 ml glass tubes containing a solvent mixture of chloroform and methanol (1:1 v/v). The final dilution was 1:50 for fat and 1:10 for liver tissues (1 g wet material corresponds to 1 ml). Tissue samples were homogenized, and total lipids were extracted for FA analysis.

Total lipids of the homogenized liver samples were extracted using a mixture of water, chloroform and methanol (0.8:0.5:1.5 v/v/v) [21]. Next, 0.2% butylated hydroxytoluene (BHT) in methanol was added to increase the oxidative stability of lipids during the extraction procedure. Following intensive shaking, two phases were generated by the addition of a chloroform/water solution (1:1 v/v). After centrifugation at 4,500 rpm for 10 min at 15 °C, the lower layer (chloroform phase) was collected, chloroform was evaporated under a gentle nitrogen stream, and the lipids were solved in a chloroform/methanol (1:1 v/v) mixture.

The different lipid classes were separated by preparative thin-layer chromatography on mm silica PSC plates (5 cm×5 cm, Merck, Darmstadt, Germany). The diluted samples, containing approximately 2.5 mg lipid, and a standard mixture consisting entirely of 1 mg/mol TAG, non-esterified FAs (NEFAs), cholesterol (C), cholesterol ester (CE) and PLs were spotted on pre-washed PSC plates. In the second step, plates were incubated in a solvent system containing chloroform and methanol (95:5 v/v) for band visualization. Single bands corresponding to the different lipid classes were identified via the authentic standards after primuline staining, scraped off and subsequently extracted and esterified for analysing FA profiles with internal standards.

A one-step lipid extraction, methylation and trans-esterification method with subsequent gas chromatography (GC) [22] was used to determine the total FA content and lipid class FA compositions of the various ATs and liver tissues, with L-phosphadidylcholin-C17:0 (0.8 mg/ml) as an internal standard.

Fatty acid methyl esters (FAMEs) were separated on a Varian CP 3800 gas chromatograph (GC, Varian, Darmstadt, Germany) equipped with a 30 m Omegawax TM 320 capillary column (0.32-mm ID, 0.2 μm df) (Supelco, Bellefonte, PA, USA). The GC oven temperature was set at 200 °C, and helium was used as a carrier gas with a flow rate of 1 ml/min. Chromatographic peaks were integrated by the Star 5.0 software (Varian) using the internal standard as the reference peak. FAMEs were used to identify and evaluate FAs assuming a direct relationship between peak area and FAME weight. Total FA was expressed as μmol/g tissue or μmol/ml, and FA composition was expressed as percent of total FA content. Blank values were subtracted in case of unavoidable contamination of reagents and solvents with some FAs (C10:0, C14:0, C16:1n7, C18:0, C22:2n6, C24:1n9) [22].

### Desaturase and elongase activity indices

Desaturase and elongase activity indices were calculated using the product/precursor ratio of the percentages of individual FAs according to the following notation: C16:1n-7/C16:0 = Δ9-desaturase (SCD16), C18:1n-9/C18:0 = Δ9-desaturase (SCD18), C18:3n-6/C18:2n-6 = Δ6-desaturase (D6D), and C20:4n-6/C20:3n-6 = Δ5-desaturase (D5D) and C18:0/C16:0 = elongase (Elo).

### Statistical analysis

Statistical analyses were performed using a statistical software program (Statistica, StatSoft GmbH, Hamburg, Germany). Data were tested for normality by the Shapiro-Wilk test. Levene’s test was used to assess equality of variance. A non-parametric Mann-Whitney U-test was applied to determine differences in the lipid FA composition and FA ratios between horses and ponies. The Kruskal-Wallis ANOVA was used to compare different locations for FA distribution and FA ratios. A two-tailed Dunn’s test correcting for multiple comparisons was done as a post hoc test. Differences were considered significant at *P* values lower than 0.05. Data are presented as medians with 25^th^ and 75^th^ percentiles.

## Results

### FA composition of adipose tissue

The total lipid FA concentrations of RPN-, MSC- and SCfat were similar between horses and ponies, but higher FA levels were found in the liver tissue of ponies than in those of horses (Table 1). Saturated fatty acids (SFAs) and n-9 monounsaturated fatty acids (MUFAs) were the most abundant FA groups in the RPN-, the MSC- and SCfat depots (Table 2) in both horses and ponies. There were no significant differences in the percent composition of the SFA, MUFA and iso-branched-chain FA (iso-BCFA) fractions between horses and ponies. Horses had a significantly higher amount of n-6 PUFAs (*P* < 0.01) and a trend for lower n-3 PUFAs than ponies.

**Table 1.**
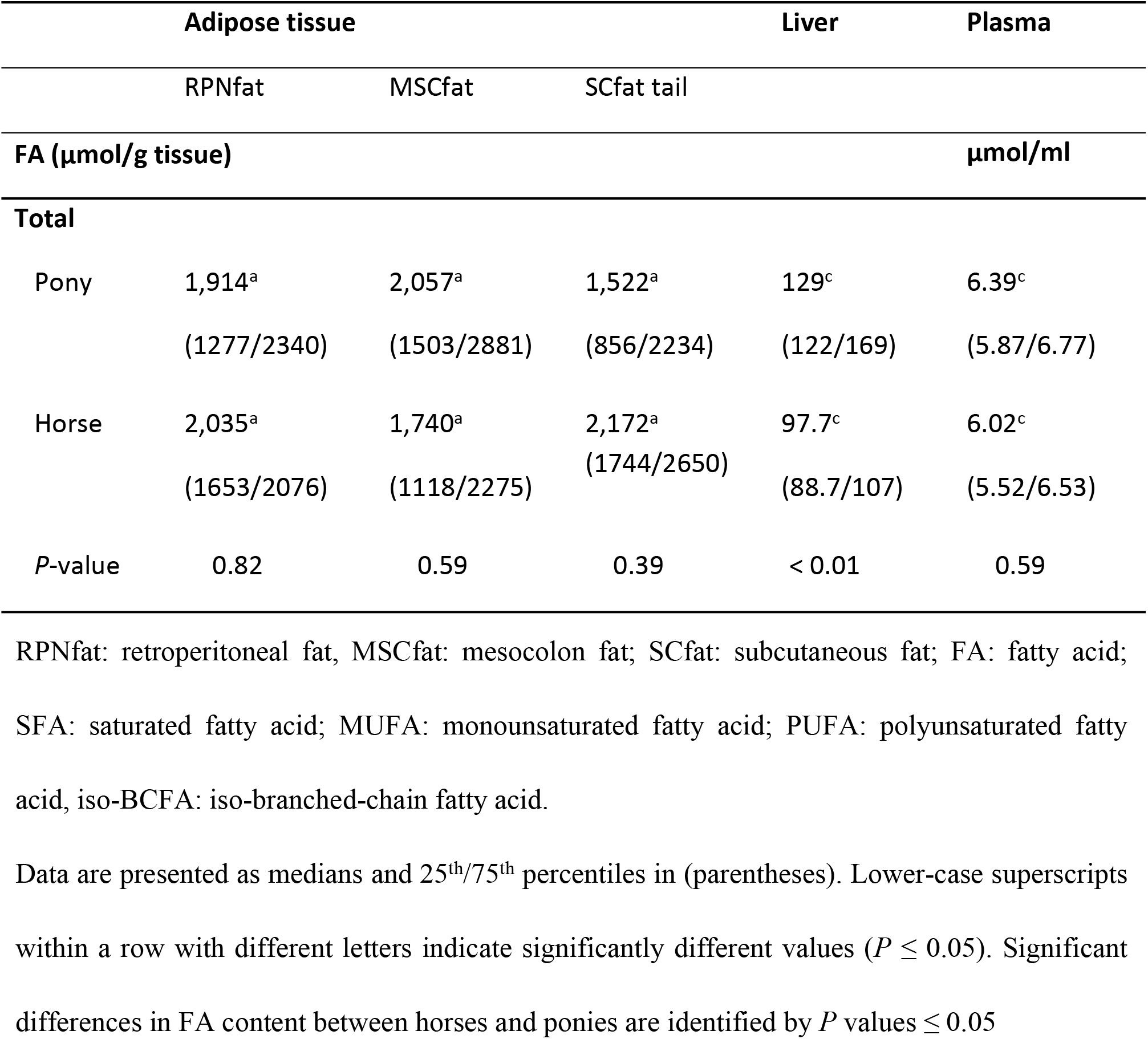
FA concentrations of different ATs, liver (μmol/g tissue) and plasma (μmol/ ml) in horses (n=6) and ponies (n=6).

**Table 2.** FA composition (% of total FA amounts) of the different ATs, liver and plasma in horses (n=6) and ponies (n=6).

|  | **Adipose tissue** |  |  | **Liver** | **Plasma** |
| --- | --- | --- | --- | --- | --- |
|  | RPNfat | MSCfat | SCfat tail |  |  |
| **FA (%) of total FA** |  |  |  |  |  |
| **SFA** |  |  |  |  |  |
| Pony | 37.9<sup>a</sup> | 34.5<sup>ab</sup> | 36.0<sup>ab</sup> | 39.5<sup>a</sup> | 31.1<sup>b</sup> |
|  | (36.0/41.3) | (32.8/35.1) | (32.3/38.2) | (38.2/40.7) | (30.6/31.5) |
| Horse | 40.1<sup>a</sup> | 37.1<sup>ab</sup> | 35.9<sup>ab</sup> | 43.1<sup>a</sup> | 30.8<sup>b</sup> |
|  | (36.5/41.3) | (35.1/38.6) | (34.7/37.4) | (39.9/47.8) | (30.7/30.9) |
| *P*-value | 0.48 | 0.09 | 0.94 | 0.13 | 0.67 |
| **MUFA** |  |  |  |  |  |
| **n-7** |  |  |  |  |  |
| Pony | 6.42<sup>abc</sup> | 7.68<sup>ac</sup> | 8.41<sup>a</sup> | 3.06<sup>b</sup> | 3.16<sup>bc</sup> |
|  | (6.08/10.0) | (6.58/10.0) | (7.89/9.06) | (2.67/3.94) | (2.42/4.09) |
| Horse | 6.08<sup>abc</sup> | 6.27<sup>ab</sup> | 8.38<sup>a</sup> | 2.49<sup>c</sup> | 2.63<sup>bc</sup> |
|  | (5.34/6.98) | (5.75/7.05) | (6.55/9.30) | (2.34/2.64) | (2.47/2.66) |
| *P*-value | 0.39 | 0.13 | 0.94 | 0.09 | 0.39 |
| **n-9** |  |  |  |  |  |
| Pony | 27.6<sup>ab</sup> | 31.1<sup>a</sup> | 29.6<sup>a</sup> | 13.7<sup>b</sup> | 13.9<sup>b</sup> |
|  | (25.5/32.3) | 26.9/34.3) | (29.1/33.4) | (12.7/17.5) | (13.4/15.7) |
| Horse | 28.4<sup>ab</sup> | 30.4<sup>a</sup> | 30.7<sup>a</sup> | 11.7<sup>b</sup> | 12.9<sup>b</sup> |
|  | (27.8/29.0) | (29.9/31.1) | (29.4/33.0) | (10.9/11.7) | (12.3/14.4) |
| *P*-value | 0.59 | 0.70 | 0.70 | 0.03 | 0.18 |
| n-11 |  |  |  |  |  |
| Pony | 0.07<sup>a</sup> | 0.06<sup>a</sup> | 0.07<sup>a</sup> | 0.15<sup>b</sup> | 0.10<sup>ab</sup> |
|  | (0.06/0.09) | (0.06/0.07) | (0.06/0.07) | (0.11/0.19) | (0.09/0.14) |
| Horse | 0.07<sup>ab</sup> | 0.08<sup>ab</sup> | 0.06<sup>ab</sup> | 0.02<sup>a</sup> | 0.09<sup>b</sup> |
|  | (0.06/0.08) | (0.06/0.10) | (0.05/0.08) | (0.02/0.02) | (0.09/0.11) |
| *P*-value | 1.00 | 0.24 | 0.82 | < 0.01 | 0.94 |
| **PUFA** |  |  |  |  |  |
| n-3 |  |  |  |  |  |
| Pony | 17.0<sup>a</sup> | 17.4<sup>a</sup> | 16.8<sup>a</sup> | 10.1<sup>ab</sup> | 6.42<sup>b</sup> |
|  | (13.2/19.0) | (12.6/21.6) | (11.1/19.4) | (7.38/11.9) | (5.80/7.05) |
| Horse | 12.3<sup>a</sup> | 12.0<sup>a</sup> | 12.0<sup>a</sup> | 8.03<sup>ab</sup> | 4.27<sup>b</sup> |
|  | (10.8/15.0) | (11.5/14.7) | (10.3/15.0) | (7.07/8.9) | (3.88/4.51) |
| *P*-value | 0.09 | 0.13 | 0.24 | 0.24 | 0.02 |
| n-6 |  |  |  |  |  |
| Pony | 6.77<sup>a</sup> | 7.03<sup>a</sup> | 6.71<sup>a</sup> | 22.4<sup>ab</sup> | 41.8<sup>b</sup> |
|  | (6.34/7.82) | (6.34/7.61) | (5.93/7.77) | (21.4/23.1) | (40.9/46.0) |
| Horse | 10.9<sup>ab</sup> | 10.6<sup>ab</sup> | 9.12<sup>a</sup> | 30.0<sup>bc</sup> | 47.6<sup>c</sup> |
|  | (9.83/14.4) | (10.55/13.9) | (8.70/12.4) | (27.8/37.1) | (46.3/48.7) |
| *P*-value | < 0.01 | < 0.01 | < 0.01 | < 0.01 | 0.02 |
| **Iso-BCFA** |  |  |  |  |  |
| Pony | 0.47<sup>a</sup><br>(0.30/0.73) | 0.48<sup>a</sup><br>(0.30/0.78) | 0.53<sup>a</sup><br>(0.35/0.75) | 10.0<sup>b</sup><br>(9.57/10.3) | 0.66<sup>ab</sup><br>(0.62/0.89) |
| Horse | 0.51<sup>abc</sup><br>(0.46/0.57) | 0.50<sup>ab</sup><br>(0.44/0.60) | 0.50<sup>a</sup><br>(0.42/0.55) | 1.89<sup>c</sup><br>(1.38/2.26) | 0.71<sup>bc</sup><br>(0.60/0.80) |
| *P*-value | 0.70 | 1.00 | 0.82 | < 0.00 | 0.82 |
| **n-6/n-3** |  |  |  |  |  |
| Pony | 0.40<sup>a</sup><br>(0.33/0.59) | 0.40<sup>a</sup><br>(0.29/0.60) | 0.40<sup>a</sup><br>(0.31/0.69) | 2.22<sup>ab</sup><br>(1.80/3.13) | 6.50<sup>b</sup><br>(5.80/7.93) |
| Horse | 0.89<sup>a</sup><br>(0.66/1.33) | 0.88<sup>a</sup><br>(0.71/0.95) | 0.76<sup>a</sup><br>(0.58/1.20) | 3.74<sup>ab</sup><br>(3.12/5.25) | 11.2<sup>b</sup><br>(10.3/12.55) |
| *P*-value | < 0.01 | < 0.01 | 0.02 | < 0.01 | 0.02 |

RPNfat: retroperitoneal fat, MSCfat: mesocolon fat; SCfat: subcutaneous fat; FA: fatty acid; SFA: saturated fatty acid; MUFA: monounsaturated fatty acid; PUFA: polyunsaturated fatty acid, iso-BCFA: iso-branched-chain fatty acid.
Data are presented as medians and 25<sup>th</sup>/75<sup>th</sup> percentiles in (parentheses). Lower-case superscripts within a row with different letters indicate significantly different values ( $P \leq 0.05$ ). Significant differences in FA composition and FA ratio between horses and ponies are identified by $P$ values $\leq 0.05$

### FA composition of the liver

In addition to SFAs, PUFAs of the n-6 FA family were the most dominant lipid FA in the liver (Table 2). The hepatic n-3 PUFA content was not different between ponies and horses, but the percentage of the hepatic n-6 PUFA fraction was significantly higher in horses than in ponies. The n-6/n-3 ratios calculated for all tissues and plasma in horses were significantly lower than those in ponies. Except for the total hepatic n-11 MUFA fraction in ponies, MUFA contents were lower in the liver than in AT depots for both horses and ponies. The total n-11 MUFA concentration in the liver of ponies was 5-fold higher than the corresponding MUFA content in horses.

Ponies contained a 2-fold higher proportion of n-11 MUFAs in the liver than in the different ATs. Liver iso-BCFA content (10.0%) was 5-fold higher in ponies than in horses.

### FA composition of plasma

In horses and ponies, SFAs and PUFAs of the n-6 FA family represented the majority of plasma lipids (Table 2).

### Tissue comparison

A comparison of the different fat depots with liver tissue showed that horses and ponies have inverted n-6/n-3 PUFA ratios and higher percentages of iso-BCFAs in the liver than in the AT depots (Table 2).

Δ9-desaturase activity indices determined from the 16:1n7/16:0 ratio and 18:1n9/18:0 ratio were significantly lower in the liver than in the ATs for both horses and ponies (Fig 1A and B). Increased Δ6- and Δ5-desaturase and elongase indices were found in the liver compared to the AT depots in all animals (Fig 1C-E). In this context, horses had a significantly higher hepatic Δ5-desaturase index than ponies (Fig 1D).

**Fig 1.**
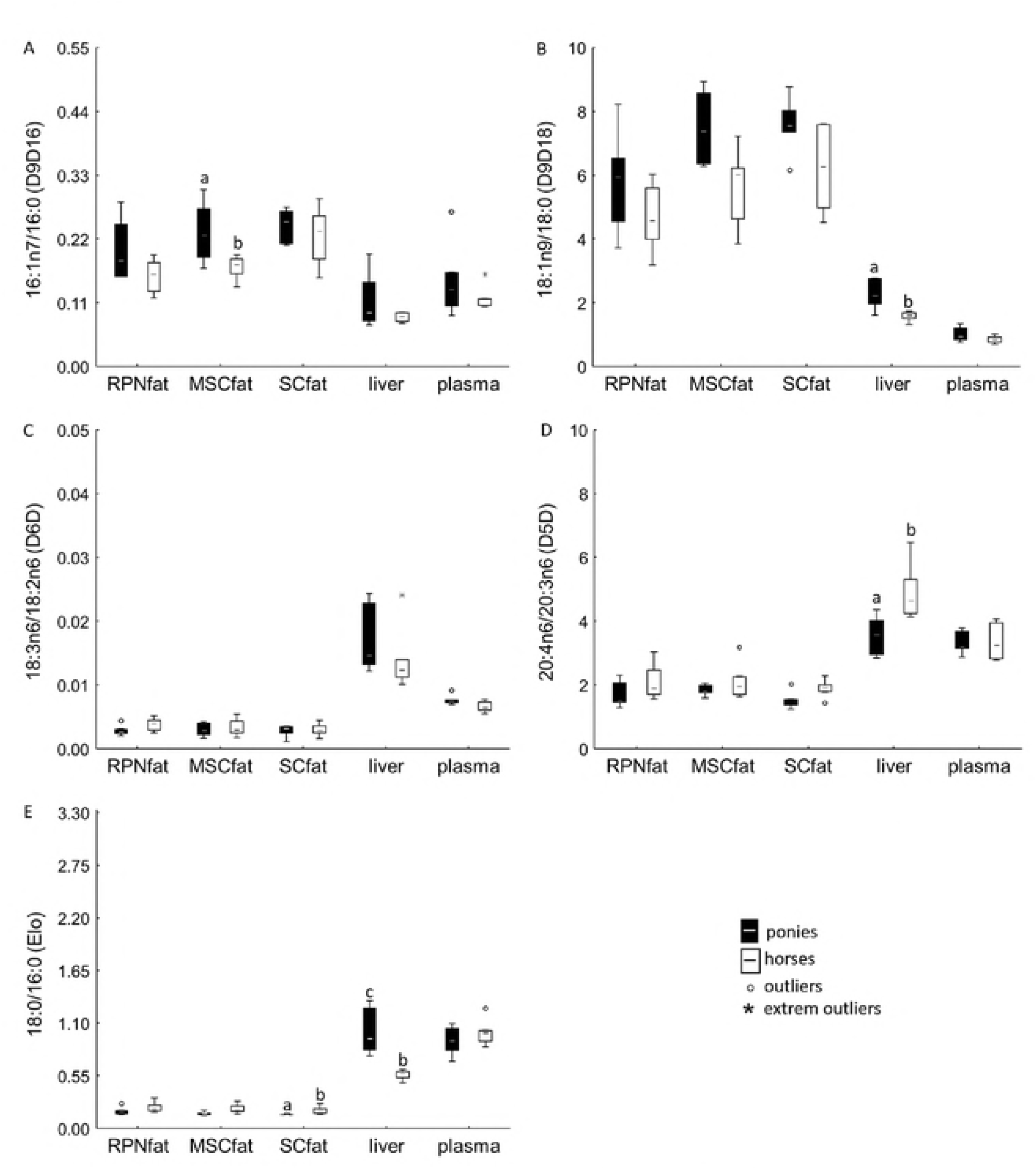
Calculated Desaturase and Elongase Activity Indices. Product/precursor ratio of the percentages of individual FAs represent desaturase and elongase activity indices: (A) C16:1n-7/C16:0 = Δ9-desaturase (D9D16), (B) C18:1n-9/C18:0 = Δ9-desaturase (D9D18), (C) 18:3n-6/C18:2n-6 = Δ6-desaturase (D6D), (D) C20:4n-6/C20:3n-6 = Δ5-desaturase (D5D) and (E) C18:0/C16:0 = elongase (Elo). Data are shown as whisker plots. Boxes represent the interquartile range (IQR) between the 25^th^ and 75^th^ percentiles. Horizontal lines are medians. Error bars show the full range excluding outliers (dots), which are defined as being more than 1.5 IQR outside the box. Lower-case superscripts with different letters indicate significant differences in these values between equine species (*P* ≤ 0.05).

Plasma FA lipid composition (Table 1b) and Δ9-, Δ6- and Δ5-desaturase and elongase indices corresponded more closely to the liver profile than to the AT profiles in both horses and ponies (Fig 1A-E).

In all animals, the majority of hepatic FAs accumulated in the polar PL fraction. Among the neutral FA fractions, TAGs had the highest FA amount, followed by NEFAs and CEs (Table 3). The absolute amounts of FAs in the PL fraction were significantly higher in ponies than in horses (*P* = 0.004). In ponies, the FA levels in the hepatic NEFA fraction were 2-fold higher (*P* = 0.03) and that of the CE fraction 3-fold higher (*P* = 0.004) than those in horses. There were no significant differences in the FA levels of TAGs except for one pony that indicated TAG FA values 10-fold higher than the average FA content measured in the TAG fractions of the other animals.

**Table 3.** FA content (μmol/g) of the separated hepatic lipid classes in horses (n=6) and in ponies (n=6).

|  | PL | NEFA | TAG | CE |
| --- | --- | --- | --- | --- |
| FA (μmol/g tissue) |  |  |  |  |
| **Total** |  |  |  |  |
| Ponies | 94.0<sup>a</sup><br>(88.6/100) | 5.06<sup>bc</sup><br>(3.52/6.72) | 27.9<sup>ab</sup><br>(18.9/58.7) | 2.45<sup>c</sup><br>(1.83/2.46) |
| Horses | 68.2<sup>a</sup><br>(61.4/70.7) | 2.62<sup>bc</sup><br>(2.04/2.94) | 26.2<sup>ab</sup><br>(24.5/30.2) | 0.61<sup>c</sup><br>(0.51/0.83) |
| *P*-value | < 0.01 | 0.03 | 0.82 | < 0.01 |

PL: phospholipid; NEFA: non-esterified free fatty acid; TAG: triacylglyceride; CE: cholesterol ester; RPNfat: retroperitoneal fat, MSCfat: mesocolon fat; SCfat: subcutaneous fat; FA: fatty acid; SFA: saturated fatty acid; MUFA: monounsaturated fatty acid; PUFA: polyunsaturated fatty acid, iso-BCFA iso-branched-chain fatty acid.
Lower-case superscripts within a row with different letters indicate significantly different values ( $P \leq 0.05$ ). Significant differences in FA content between horses and ponies are identified by *P* values $\leq 0.05$ .

N-6 PUFAs were the most abundant FAs in the PL fractions of horses and ponies, followed by SFAs, n-9 MUFAs, n-3 PUFAs together with iso-BCFAs and, at last, n-7 and n-11 MUFAs (Table 4). Compared to horses, ponies had significant higher n-9 MUFA and n-3 PUFA levels but significant lower n-6 PUFA contents in the PL fraction, resulting in a lower n-6/n-3 ratio. No significant differences in SFA and iso-BCFA levels were found in the hepatic PL fractions between horses and ponies. Similar to the PL fraction, n-6 PUFA levels and calculated n-6/n-3 ratios in NEFAs, TAGs and CE were significantly higher in horses than in ponies. N-11 MUFAs and iso-BCFAs were absent from the hepatic NEFA fraction of horses. In ponies, the hepatic CE fraction had significantly higher n-3 PUFA levels but lower SFA amounts than that in horses. Ponies had the highest iso-BCFA levels and the lowest n-9 PUFA contents in the hepatic CE fraction. These FAs were completely absent from the CE fraction of the horse liver.

**Table 4.**
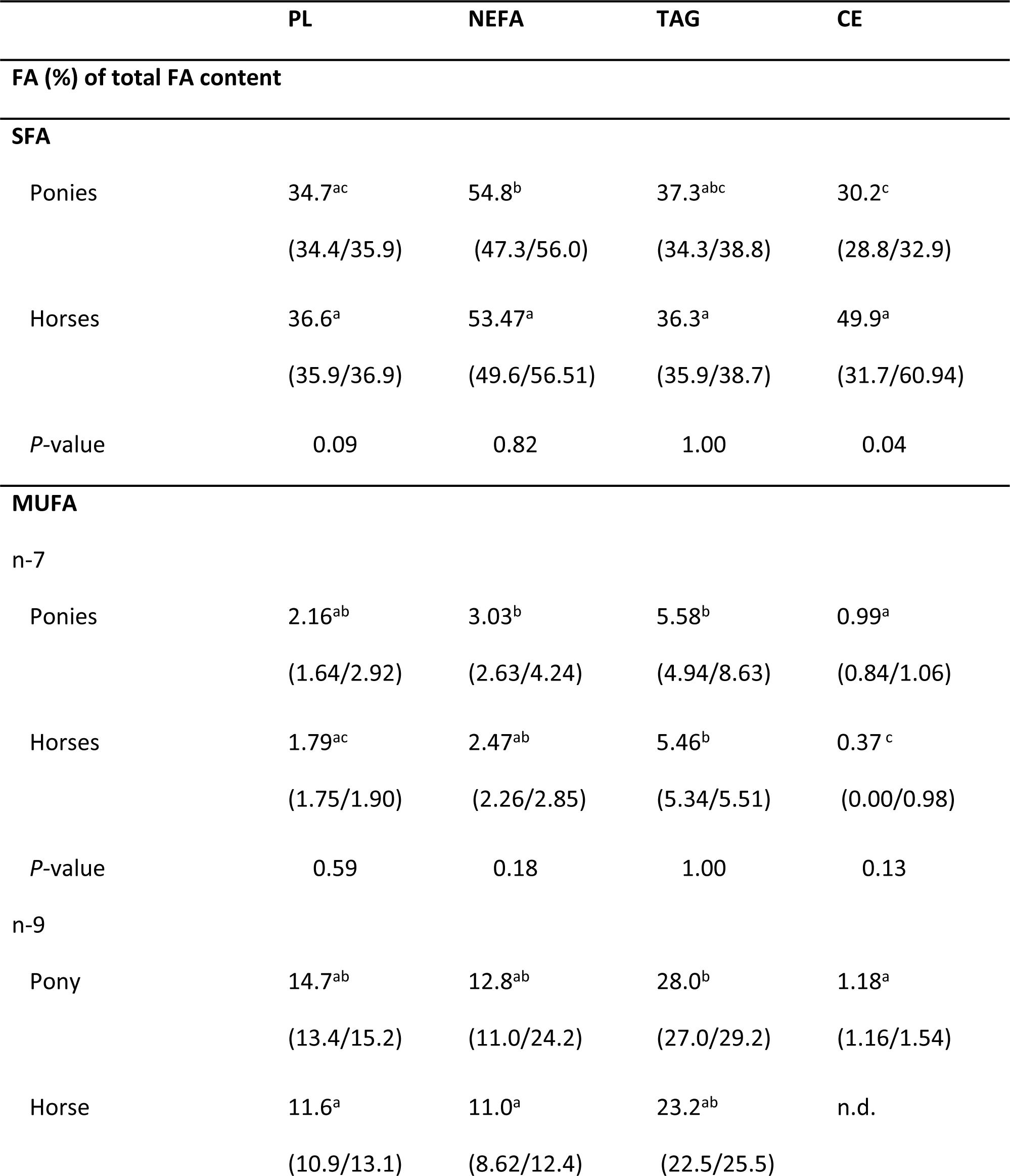
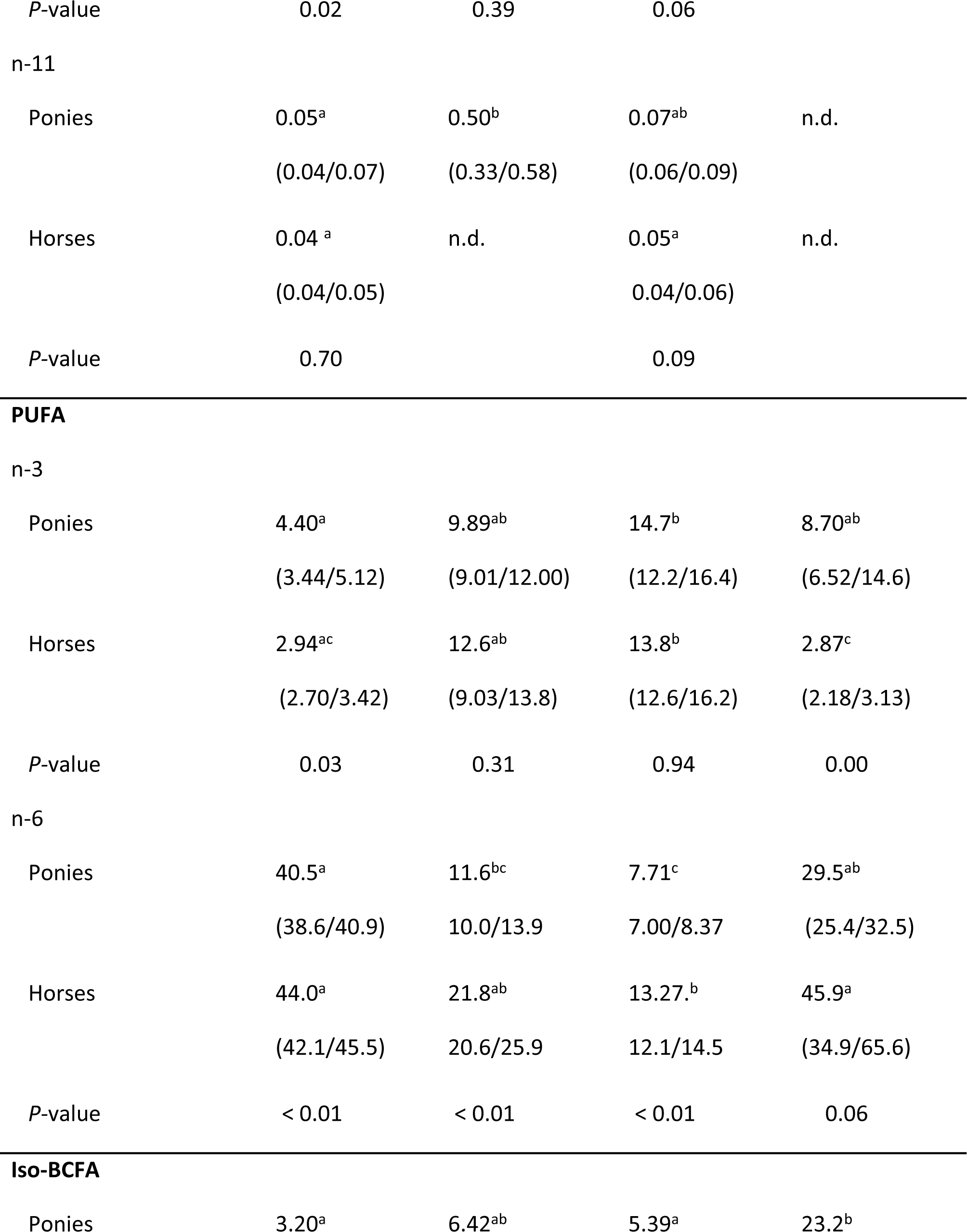
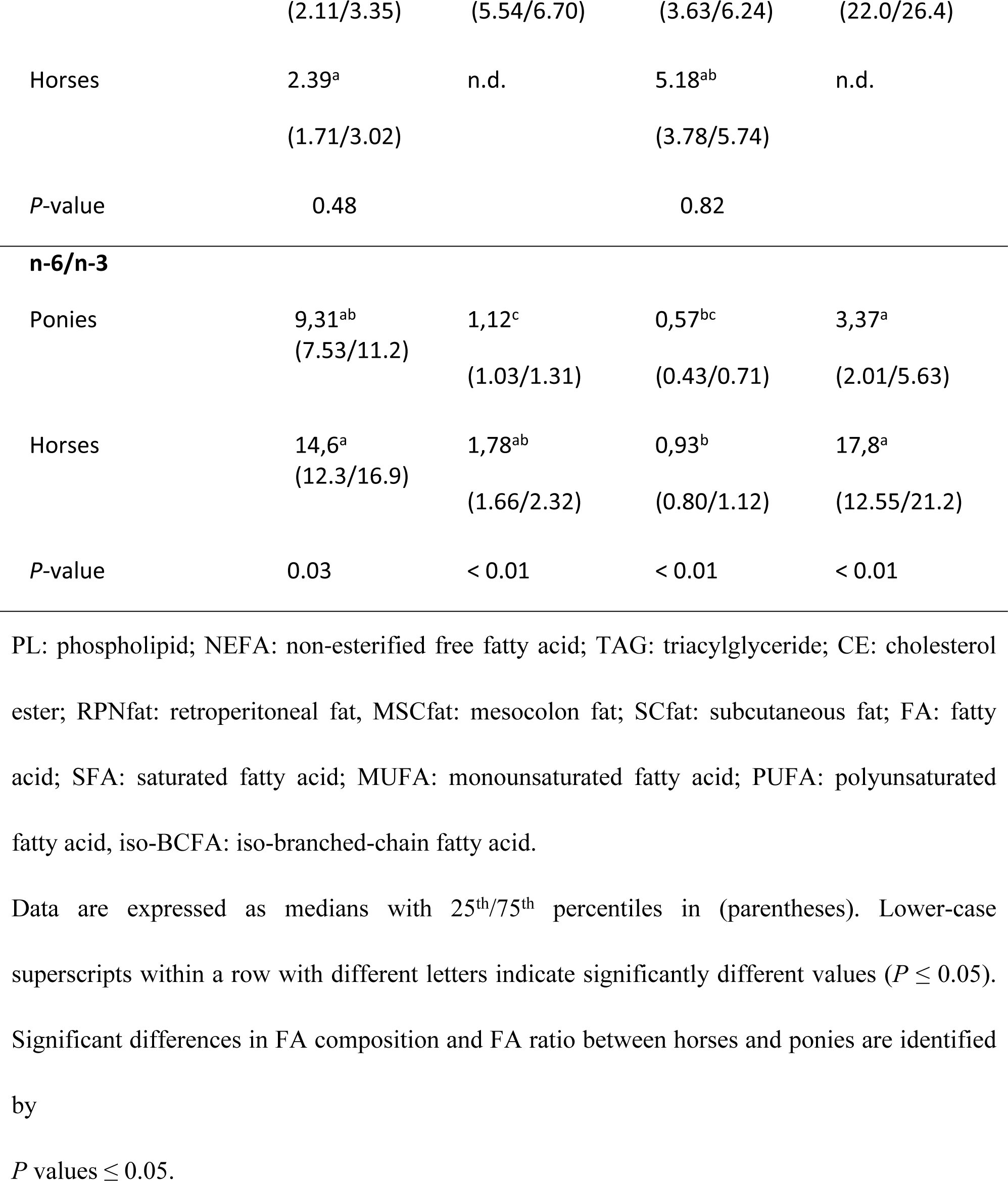
FA composition (% of total FA amounts) of the separated hepatic lipid classes in horses (n=6) and in ponies (n=6).

## Discussion

The current study aimed to compare FA profiles of ATs, liver and plasma between lean Shetland ponies and lean Warmblood horses. For general background information, healthy animals with a moderate body condition score (BCS) [18] and cresty neck score (CNS) [23] were included in the ongoing study. Using standardized conditions, our study provides key information on the differences in FA class distribution of lipids in different tissues by comparing lean ponies with lean horses under controlled feeding and management conditions.

The regulating impact of ATs on lipid metabolism and tissue crosstalk seemed to be different in horses and ponies [24]. Ex vivo studies on the lipolysis activity of equine adipocytes highlighted a disturbed NEFA release from TAG fat stores as a cause for hyperlipidaemia in ponies but not in horses under negative energy intake [24]. However, little is known about the function of the FA lipid profile in AT metabolism in equids, although FAs are known to act as regulatory key players in various physiological, metabolic and inflammatory processes [7,25].

In equids, most studies on the AT lipid pattern have mainly focused on the intramuscular fat depots [26-30]. To our knowledge, this study is the first to investigate differences in the FA composition of various visceral and subcutaneous fat depots between horses and ponies. The lipid FA composition and lipid metabolism of tissues is substantially influenced by the diet [28,31,32]. To minimize dietary effects, feeding protocols were standardized in animals by feeding them meadow hay for several weeks. In the present study, no significant differences in the total FA content and FA lipid profile between the distinct AT locations were found in horses and ponies (Table 1).

Our results confirmed previous studies that highlighted n-3 PUFAs and, to a lesser extent, n-6 PUFAs, as well as SFAs and n-9 MUFAs, as the main FA classes in ATs [28,29]. The predominance of PUFAs in ATs is well explained by the meadow hay intake [30] and the low biohydrogenation activity in the gut, resulting in an efficient uptake and deposition of PUFAs from grass species rich in n-3 and n-6 PUFAs into tissues [26,33].

In addition to ATs, the liver plays a major role in FA metabolism, which argues that researchers should focus more closely on hepatic FAs and their metabolism [34]. However, data on hepatic lipid composition and FA metabolism in equids are rare. As expected, the total FA content in liver tissue was significantly lower than that in the different ATs. In the liver, FAs mainly accumulate in the polar PL fraction, followed by the neutral fractions of TAGs, NEFAs and CEs for both horses and ponies (Table 3). Interestingly, SFA levels were quite similar between ATs and liver, but significant differences were found in MUFA and essential n-6 PUFA levels (Table 2). FAs in adipocytes mainly result from dietary FAs, which enter the adipocytes to form neutral TAGs for energy storage, thereby primary accumulating SFAs and MUFAs [35]. Furthermore, the majority of *de novo* synthesis of non-essential FAs may occur in the AT depots and not in the liver, as has been recently described for equids [36]. Data from the present study, showing significantly higher 16:1n-7/16:0 and 18:1n9/18:0 ratios in the AT depots than in liver tissue, supports these findings by reflecting a higher AT Δ9-desaturase activity (Fig 1A and B). Following *de novo* lipogenesis of saturated palmitic acid (C16:0) and stearic acid (C18:0) in the cell, ∆-9 desaturase catalyses the production of monounsaturated palmitoleic acid (C16:1n-7) from the 16:0 precursor and of oleic acid (C18:1n-9) from its 18:0 precursor [37]. Thus, the accumulation of C16:1-n7 and 18:1n9 as the main products of lipogenesis could reflect high *de novo* lipogenesis activity in AT depots.

In the liver, essential FAs are used as precursors for other FAs, especially long-chain PUFAs. In addition, FAs can be converted to biologically active FA-derived compounds such as eicosanoids that regulate a variety of physiological processes [38].

The n-6 PUFA contents being higher in the liver than in the ATs might be explained by a higher percentage of PLs being present in the membranes of the hepatocytes than in those of adipocytes. Namely, n-6 PUFA arachidonic acid (AA) is an important component of the membrane PLs. Hepatocytes preferably incorporate PUFAs, especially members of the n-6 PUFA family, in the polar PL structures of the cellular membranes [34] rather than in neutral storage lipids such as TAGs. In adipocytes, high amounts of FAs accumulate in cytosolic TAGs for energy storage, mainly including SFAs and MUFAs and smaller amounts of PUFAs [35]. Higher n-6 PUFA levels resulted in reduced n-3 PUFA levels, explaining the n-3/n-6 ratio being inverted between liver and ATs (Table 2).

Species-derived differences in the FA profile of selected ATs and liver were found for n-6 PUFAs. Horses had significantly higher n-6 PUFA contents in the ATs and liver compared to ponies. As the diets were standardized between horses and ponies, the differences in n-6 PUFAs might be related to genetically or metabolically derived differences in the incorporation, utilization or storage of PUFAs. Studies in humans confirm that in addition to nutritional influences, genetic background is highly important for PUFA composition in tissues [39]. Genetic association studies on the FA composition of ATs, serum PLs and erythrocyte membrane PLs in humans of different ethnic backgrounds revealed that polymorphisms (SNPs) in the desaturase *FADS* gene clusters [40-43] determine the efficiency of n-3 and n-6 PUFA processing, thus affecting total PUFA status. Compared to major allele carriers, minor allele carriers of the *FADS* SNPs had minor desaturase activities, indicating a high association of genotype and absolute endogenous n-3 and n-6 PUFA levels [43]. The data from the current study confirm significant differences in the desaturase and elongase activity indices between horses and ponies for liver tissue and some ATs (Fig 1A, B, D and E). This result might support the hypothesis of a variant genotype within the equids. In addition, genetic variations of other candidate genes were evidenced affecting PUFA binding, translocation and transport in human and animal tissues [39,42,44-46]. Furthermore, enzyme selectivity for specific FAs, rates of FA uptake, FA mobilization and FA reuptake were assumed to affect the PUFA composition of ATs in humans and different animal models [44-47]. In this context, FA chain length and degree of unsaturation are critical factors that might affect the individual FA supply to tissues [48]. For n-3 PUFAs, slower uptake into ATs [31] and higher mobilization [49] in relative to their values in other FAs have been observed in humans. Thus, differences in the n-6 and n-3 PUFA tissue availability in the equine sub-species might be a result of nutrigenetic interactions of dietary PUFAs and variations in genes encoding for PUFA enzymes and transporters.

In horses, the higher n-6 PUFA levels seemed surprising. PUFAs of the n-6 FA family are considered pro-inflammatory [9], which may predispose individuals to inflammatory responses and metabolic dysregulation [38,50,51]. Among n-6 PUFAs, AA in particular has signalling effects that may create a pro-inflammatory, pro-allergic and pro-tumour environment [7]. AA mainly acts as a precursor for eicosanoids and derived mediators that are associated with inflammatory diseases and immune responses. In addition, free AA may directly promote inflammatory processes by activating the transcription factor Nuclear factor kappa B (NFκB), which regulates the expression of genes associated with innate and adaptive immunity [52]. The essential linoleic acid (LA) (18:2n6), as an integral part of cellular membrane ceramides, is important in skin and barrier function [53]. LA can be metabolized to AA by several desaturase and elongase enzyme activities. Several *in vitro* studies on macrophages and *in vivo* studies in humans have indicated that LA has only a limited effect on inflammation with respect to a reduced release of pro-inflammatory cytokines such as IL-1β and IL-6 [54,55]. Considering the range of different biological effects of the two n-6 PUFAs, it seems no longer relevant to describe the functional impacts of FA families or classes; rather, the activities of individual FAs and their relevance to health and disease risk should be detailed [7]. In earlier studies on the FA composition of serum PLs in equids, it was evidenced that ponies and horses contained higher LA contents despite having less AA in serum lipids than other herbivores [1,56,57]. The inverse relationship of the two n-6 PUFAs in serum might have particular relevance in prostaglandin production. For example, human endothelial cells [58], skin fibroblasts [59] or mouse macrophages [60] enriched with LA showed reduced AA contents in the cellular PLs and reduced prostaglandin (PGI_2_) release. It is speculated that these two n-6 PUFAs also have an inverse relationship in the hepatic lipid stores of equids. Further investigations of LA and AA metabolism in relation to prostaglandin syntheses and inflammatory tissue responses in horses and ponies are needed.

Furthermore, the combined influences of n-3 and n-6 PUFAs on several regulatory and signalling systems [7,25,55] predisposes the n-6/n-3 ratio as an additional critical factor influencing health and diseases in humans and animals [50,61]. In the present study, both horses and ponies had n-6/n-3 ratios < 1 in ATs and of 2 or 3 in the liver. Similar results have been reported for subcutaneous ATs in healthy horses of different breeds [26]. Further investigations on the n-6/n-3 PUFA ratio in cases of metabolic dysregulation are important to determine optimal tissue and individual dietary ratios. The importance of the dietary n-6/n-3 ratio in diverse chronic health conditions such as obesity-linked inflammation and insulin dysregulation was evidenced in Sprague-Dawley rat models and human studies [50,61,62]. Reduction in n-6 PUFAs in the diet to a n-3/n-6 PUFA ratio of 1:1 was evidenced to prevent excessive n-6 eicosanoid action and Toll-like receptor 4 (TLR4)-induced production of pro-inflammatory cytokines via an effective blocking of corresponding signalling pathways by n-3 PUFA action. This approach avoids metabolic disorders and is beneficial for health. Use of n-6 PUFA-dominated dietary ratios of 1:4 failed to result in these changes [61,62]. However, it cannot be excluded that the adaptation period to the same diet was too short to exclude any dietary influences on PUFAs in the present study.

Another interesting finding of the current study was the high iso-BCFA content in the liver of horses and ponies compared to that in the ATs (Table 2). In mammals, BCFAs are found in several tissues, including skin, brain, blood and cancer cells [63-66]. According to their structure and main metabolic origin, several types of BCFAs were differentiated. Iso- and anteiso-BCFAs, also called terminal BCFAs, are the main components of bacterial membranes and are synthesized from branched-chain amino acids and their corresponding branched-short-chain carboxylic acids [67]. However, the non-terminal BCFAs were reported to be synthesized in tissues [68]. Although the function and physiological roles of BCFAs are rarely understood, their wide distribution in different tissues might suggest an important function in several metabolic processes. Studies in humans reported anti-inflammatory and insulin-synthesizing effects of BCFAs [63,69]. It is speculated that BCFAs may affect metabolic genes and transcription factors [64].

However, little information is available on the role of BCFAs in horses. Santos et al. [70] found high levels of BCFAs in the equine hindgut, which might be related to bacterial and protozoan mass or bacterial and protozoan fermentation. It is well known from ruminants that the bacterial BCFAs of the rumen can be absorbed and incorporated into TAG tissues, thus increasing the terminal BCFA content in ATs and milk lipids [67]. Likewise, with regard to our findings in ATs and plasma, Belaunzaran et al. [71] determined similar levels of BCFAs in horse meat and ATs and assumed that BCFA absorption occurred from diet rather than by absorption of fermentation products. Previous studies that confirm various iso- and anteiso-BCFAs as major components in plant surface waxes [72] support the idea of dietary uptake of BCFAs by tissues. Nevertheless, data on BCFA status in liver tissue were not obtained. In accordance with our data, high BCFA levels, ranging from 2.1% to 2.5% of total FA content, were found in the liver of ruminants [67]. The high hepatic BCFA amounts in bovines were thought to be associated with TAG accumulation. This link was also confirmed in the current study. However, the highest iso-BCFA contents were highlighted in the CE fraction of the pony liver, in contrast to a complete lack of hepatic CE iso-BCFAs in horses. Clearly, there exists species-derived differences in iso-BCFA metabolism that might be linked to cholesterol metabolism in equids. We further speculate that different expression patterns of receptors and enzymes are involved in intracellular degradation, turnover and transport of iso-BCFAs through the plasma membranes in ponies and horses. Further research on the entire role and metabolism of iso-BCFAs in hepatic FA metabolism is required. In that context, it seems appropriate to investigate the function of different iso-BCFAs in context with metabolic dysregulation in horses and ponies.

Interestingly, horses rather than ponies lack some of the FA classes in different hepatic lipid fractions. Studies by Yamamoto et al. [73] confirm the lack of hepatic n-9 and n-11 MUFAs in the polar CE fraction and found comparable levels of SFAs in the CE (61.7% of total FA) and neutral TAG fraction (44.1% of total FA) as we did. However, data on the long chain FA were missing, which is partly offset by data from the current study.

FA levels and lipid profiles were similar between liver and plasma, suggesting plasma as a potential biomarker for liver lipid content and FA lipid composition [74]. These findings are further supported by the close relationship of the FA elongase and desaturase indices in the liver and plasma, which were both different from those in the ATs (Fig 1A-E).

## Conclusion

The present results provide basic data on the FA profiles in different ATs, liver and plasma in lean horses and ponies under standard conditions. The higher n-6/n-3 ratios in the ATs, liver and plasma of horses in comparison to ponies and differences in the desaturase and elongase indices may reflect breed-related differences in the acquisition (e.g., FA uptake and *de novo* lipogenesis), removal (i.e., mitochondrial FA oxidation and FA mobilization) and turnover of FA. In liver tissue, horses also had lower hepatic iso-BCFA levels than ponies. As iso-BCFAs are linked to anti-inflammatory and insulin-synthesizing effects, these findings need further elucidation.

## Acknowledgments

The authors are grateful to Gabriele Dobeleit, Sabine Klemann and Jana Tietke for their enormous technical support.

## Supporting information

**S1 Table. Descriptive data of equines.** * Mean ± SD. ** Median (25^th^/75^th^ percentile).

## Footnotes

1 Mean median of two independent evaluators.

